# Refining SARS-CoV-2 Intra-host Variation by Leveraging Large-scale Sequencing Data

**DOI:** 10.1101/2024.04.26.591384

**Authors:** Fatima Mostefai, Jean-Christophe Grenier, Raphäel Poujol, Julie G. Hussin

## Abstract

Understanding the evolution of viral genomes is essential for elucidating how viruses adapt and change over time. Analyzing intra-host single nucleotide variants (iSNVs) provides key insights into the mechanisms driving the emergence of new viral lineages, which are crucial for predicting and mitigating future viral threats. Despite the potential of next-generation sequencing (NGS) to capture these iSNVs, the process is fraught with challenges, particularly the risk of capturing sequencing artifacts that may result in false iSNVs. To tackle this issue, we developed a workflow designed to enhance the reliability of iSNV detection in large heterogeneous collections of NGS libraries. We use over 130,000 publicly available SARS-CoV-2 NGS libraries to show how our comprehensive workflow effectively distinguishes emerging viral mutations from sequencing errors. This approach incorporates rigorous bioinformatics protocols, stringent quality control metrics, and innovative usage of dimensionality reduction methods to generate representations of this high-dimensional dataset. We identified and mitigated batch effects linked to specific sequencing centers around the world and introduced quality control metrics that consider strand coverage imbalance, enhancing iSNV reliability. Additionally, we pioneer the application of the PHATE visualization approach to genomic data and introduce a methodology that quantifies how related groups of data points are within a two-dimensional space, enhancing our ability to explain clustering patterns based on their shared genetic characteristics. Our workflow sheds light on the complexities of viral genomic analysis with state-of-the-art sequencing technologies and advances the detection of accurate intra-host mutations, opening the door for an enhanced understanding of viral adaptation mechanisms.

## 1 Introduction

The advancements in high-throughput sequencing technologies have revolutionized the study of viral genomes, particularly evident in the case of SARS-CoV-2 during the COVID-19 pandemic. The ability to track the virus’s mutations and evolution during host infection is critical in understanding the emergence of various variants of concern (VOCs). These VOCs, resulting from the accumulation of mutations, demonstrate the importance of selective pressures both within an individual host (intra-host) and during transmission between hosts (inter-host) (Lauring 2020). This complex interplay is key to the evolution of viral lineages, influenced by factors like error-prone replications and host RNA-editing mechanisms (Di Giorgio et al. 2020). In the current literature, there are several hypotheses to explain the interplay between intra-host and inter-host dynamics in the development of SARS-CoV-2 VOCs (Markov et al. 2023). These hypotheses include evolution within chronically infected individuals (Sonnleitner et al. 2022; Quaranta et al. 2022; Hill et al. 2022; Ghafari et al. 2022; Oude Munnink et al. 2021; Hale et al. 2022; Oreshkova et al. 2020; Bashor et al. 2021), spillovers from animal populations (Washburne et al. 2022; Sacchetto et al. 2021; Robinson et al. 2023; Goldberg et al. 2023; Rajendran et al. 2022), and emergence in regions with limited genomic surveillance. Understanding these processes is vital to explain the rapid evolution of VOCs such as Delta and Omicron, which have shown significant evolutionary leaps.

In response to the pandemic, a vast number of next-generation sequencing (NGS) libraries for SARS-CoV-2 have been generated, primarily to construct consensus sequences for tracking inter-host mutations and VOCs. However, they also provide valuable insights into intrahost diversity, enabling the identification of intra-host single nucleotide variants (iSNVs) that are key in exploring hypotheses around VOC emergence. Despite the considerable size and breadth of available NGS libraries, the existing body of research on iSNV analysis remains limited, with the majority of studies focusing on a relatively small number of NGS libraries (Sun et al. 2023; Messali et al. 2023; Xi et al. 2023; Sun et al. 2023; Armero et al. 2021; Wertheim et al. 2022; Y. Wang et al. 2021; Sonnleitner et al. 2022; Zhang et al. 2022; Quaranta et al. 2022). This gap in research can also be attributed to challenges related to data quality, such as the presence of sequencing artifacts that introduce errors and lead to false iSNVs. To mitigate these challenges, current practices in intra-host viral analysis include the use of technical replicates (Zhang et al. 2022), which, while effective, are resource-intensive, and the application of hard filters on coverage and frequency, which lack uniformity across different studies and often overlook noteworthy sequencing artifacts like strand bias (Roder et al. 2023; Armero et al. 2021; Hedskog et al. 2010; Bull et al. 2011; Tonkin-Hill et al. 2021). This concern underscores the need for more standardized methodologies in processing complex sequencing data to ensure accurate and reliable iSNV analysis.

In computational biology, dimensionality reduction techniques are commonly used to simplify the representation and analysis of complex datasets, like viral sequencing data, and to uncover inherent data biases. These techniques, which have seen significant improvements with the rise of high-dimensional data, include Principal Component Analysis (PCA) (Novembre et al. 2008), often used for summarizing human genetic data, t-SNE (Platzer 2013; Tamazian et al. 2022) for analyzing local structures, and PHATE (Moon et al. 2019), a novel method that allows visualization of both global and local structures in high-dimensional data. Despite the potential of these methods, their application to the extensive SARS-CoV-2 data has been limited, often confined to analyzing consensus sequences (Hozumi et al. 2021; B. Wang et al. 2021; Mostefai et al. 2022). This gap highlights an opportunity for a broader application of the dimensionality reduction methods on viral genome data.

Here, we address this gap by using a comprehensive set of publicly available SARS-CoV-2 NGS libraries from the NCBI database, representing the pandemic’s initial years. We use a combination of bioinformatics tools, stringent quality control measures, and dimensionality reduction methods such as PHATE and t-SNE to identify intra-host mutations from sequencing artifacts. Our approach provides a workflow for analyzing SARS-CoV-2 sequencing data and establishes adapted thresholds for the 2020 and 2021 datasets. In this study, we establish a framework for rapid and precise analysis of intra-host viral data, aiming to support pandemic preparedness and response.

## 2 Results

### 2.1 Curation Pipeline Overview

While NGS data offers valuable insights into viral diversity and evolution, extracting meaningful information demands rigorous bioinformatics and representation approaches. A systematic methodology is crucial to process this data accurately, ensuring the reliability of identified iSNVs. We, therefore, propose a comprehensive workflow to extract meaningful intra-host mutations from NGS data. Our workflow is divided into two levels (Figure 1): the processing and quality control of a set of libraries (Figure 1A) and the processing and quality control of iSNVs within each library (Figure 1B).

**Figure 1:**
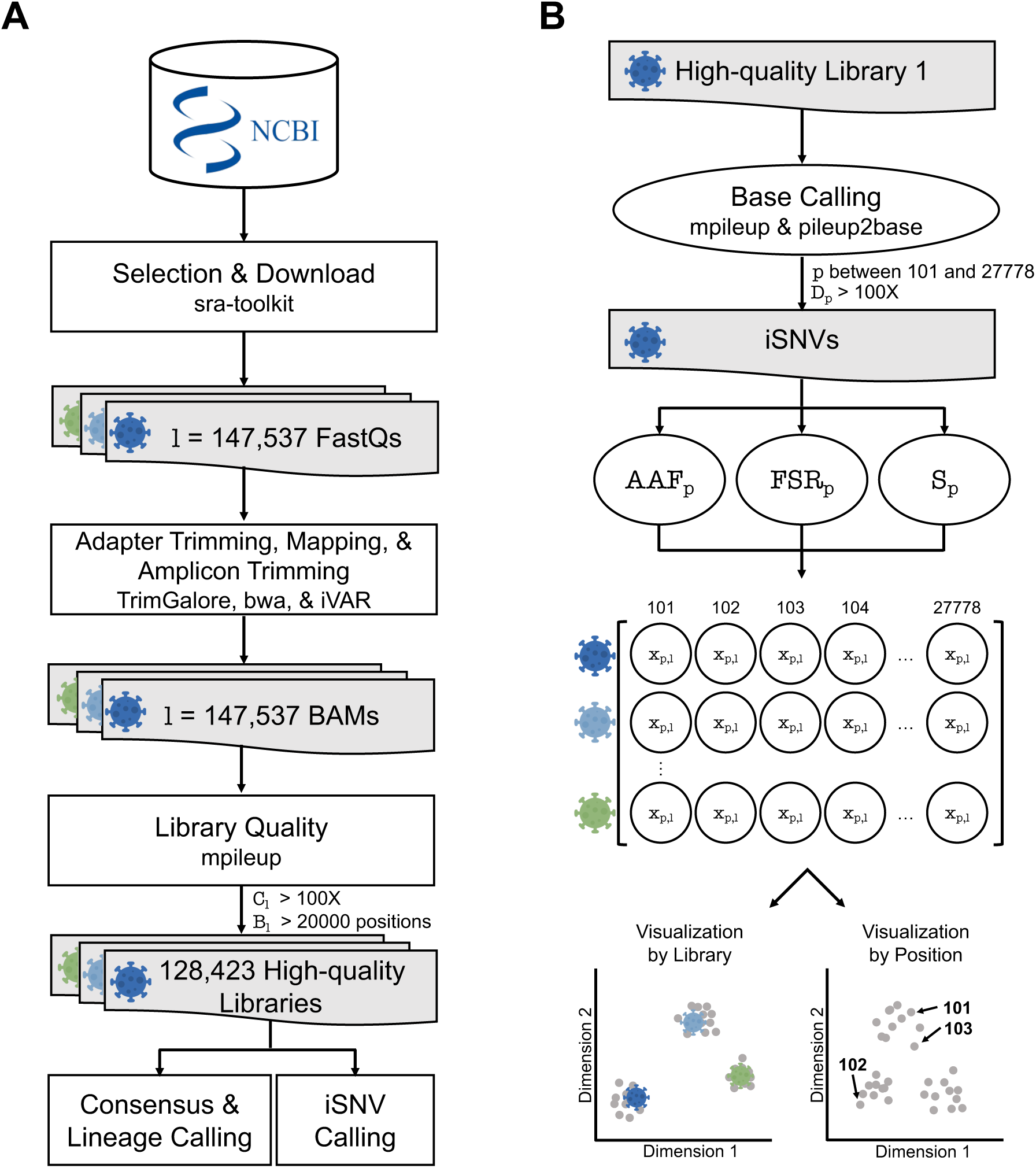
SARS-CoV-2 Sequencing Library Processing Workflows. **A**: Processing workflow of a set of SARS-CoV-2 sequencing libraries. The workflow starts with selecting and downloading 147,537 FastQ libraries from NCBI. Next, these libraries were trimmed for adapters, mapped to a reference, and trimmed again for primer targets. We set whole genome coverage filters of mean depth (*C*) *>* 100X and breadth of coverage (*B*) *>* 20000 on each library *l* to keep only high-quality libraries, keeping 128,423 libraries for further analysis. **B** Processing workflow of a single RNA sequencing library. Within each high-quality sequencing library, base calling was done to extract iSNVs. During this process, the ends of the genome were removed, keeping genomic positions *p* between 101 and 27778 and the depth *D >* 100X of *p*. Subsequently, for each iSNV, we computed the following quality metrics: Alternative Allele Frequency (*AAF*), Forward Strand Ratio (*FSR*), and Strand bias likelihood (*S*, equation 1). Thresholds for *AAF* and *S* metrics were established using dimensionality reduction visualization methods, reducing the data into two dimensions by either the libraries (left) or the genomic positions (right).

To build a set of high-quality libraries, we meticulously processed a large set of Illumina amplicon paired-end sequencing libraries, ensuring a representative sample across various time points and locations. The data processing includes adapter and quality trimming, alignment to the SARS-CoV-2 reference genome, primer trimming, and whole genome coverage quality control (see Method section 4.1). Using the processed libraries, we performed iSNV calling and computed key metrics such as Alternative Allele Frequency (*AAF*) and Strand bias likelihood (*S*) (see Methods). These metrics help to accurately identify putative iSNVs while minimizing artifacts. Next, dimensionality reduction methods, such as PHATE and t-SNE, are applied to visualize and interpret the iSNV data through analyses of clustering structures. In this process, we generate representations in two distinct ways: by the library, where each point in the visualization represents a summary of the library, and by genomic position, where each point corresponds to a specific genomic position summarizing its behaviour across libraries. PHATE maintains meaningful distance between clusters (Moon et al. 2019), preserving hierarchical relationships between sequencing libraries, we therefore use this technique to present our main results. Similar findings are observed using t-SNE (see supplementary information section 10.5). To differentiate between potential artifacts and biologically relevant patterns, the clustering structures are measured using the Percentage of Nearest Neighbors (*PNN*) presenting the same lineage label (as defined by the World Health Organization, WHO) or sequencing center (SC), providing a robust metric to quantify clustering structures of different sets of iSNVs. While the null hypothesis is for iSNVs to be randomly distributed, SC labels are used to check for associations with sequencing centers as a proxy to identify potential artifacts. Conversely, WHO labels are expected to reflect biological relevance, with the limitation that some sequencing centers favour the sequencing of some lineages over others.

### 2.2 Extracting Emerging *de novo* iSNVs

We processed 128,423 high-quality SARS-CoV-2 sequencing libraries from the first two years of the COVID-19 pandemic, ensuring a representative sampling across time and geographic locations (Figure 2A). Genome data quality was assessed by coverage depth (Figure 2B, x-axis), breadth (Figure 2B, y-axis), and strand balance (Figure 2C). The distribution of depth and breadth of coverage reveals center-specific quality variability. In turn, the strand balance coverage shows unbalanced strand coverage across the genome with an oscillating pattern at the same genomic regions independent of the sequencing center (see supplementary information section 10.2).

**Figure 2:**
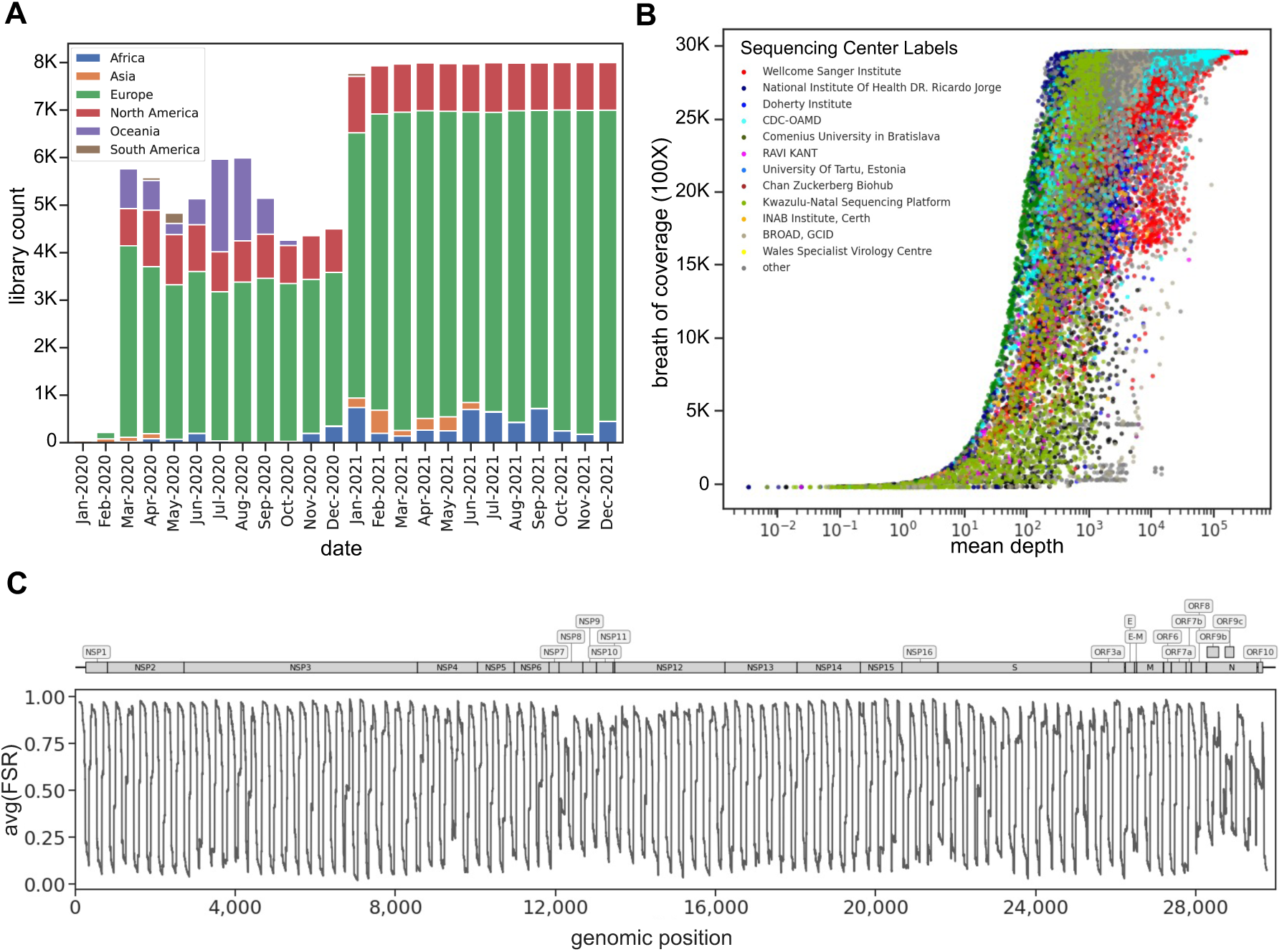
Data description and whole genome quality control. **A**: The 147,537 Illumina pairedend amplicon sequencing libraries were selected and downloaded from NCBI. Each bar in the graph represents the number of samples, categorized by their respective collection dates, with labels indicating their continents. **B**: The x-axis shows each library’s mean depth of coverage (log scale), while the y-axis shows the breadth of coverage. This breadth of coverage is the count of genomic positions covered by at least 100X of depth (*D*). Libraries with at least 1,000 libraries (93%) in our dataset are explicitly labelled with the sequencing centers, while the remaining libraries have been grouped under the “Others” label. **C**: Whole Genome forward strand ratio (FSR) averaged across libraries for each genomic position. The gene annotations are overlaid on the top panel.

We identified a total of 11,635,668 iSNVs in these libraries before any filtering steps, averaging 91 iSNVs per library (see Methods section 4.3 and Table S1). PHATE representation of the raw iSNV dataset distinctly discriminates libraries according to WHO lineage annotations (Figure 3A). The mean Percentage of Nearest Neighbours from the same WHO lineage (*PNN_W HO_*) (Figure 3B) is at 98.39%, corroborating a strong lineage-specific signature in the raw iSNVs (see Methods section 4.4).

**Figure 3:**
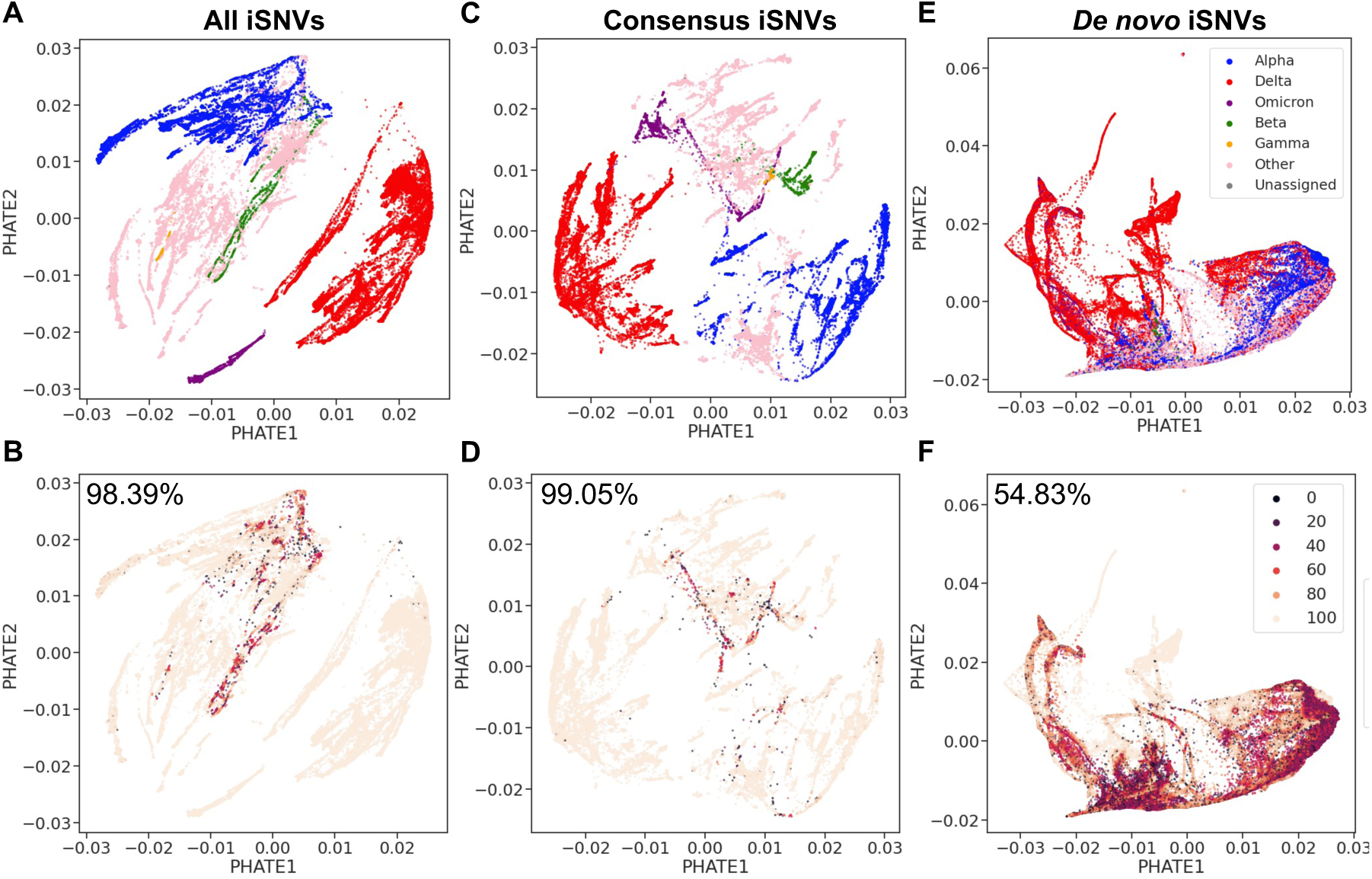
The PHATE representation organizes the unfiltered SARS-CoV-2 libraries according to WHO lineage annotation. **A**: PHATE visualization of the full dataset matrix with 11,635,668 iSNVs, using WHO lineage labels. **B**: The same dataset as **A**, labelled with the percentage of nearest neighbours that share the same WHO annotation as the library itself, and the total *PNN_W HO_* value is displayed at the top left. Darker-coloured points signify a lower percentage of neighbouring points sharing the same label as the focal point. **C**: PHATE visualization of the consensus matrix, containing 3,634,563 iSNVs, with WHO lineage labels. **D**: Consensus matrix, similar to **C**, but with *PNN_W HO_* labelling and the *PNN_W HO_* value at the top left. **E**: PHATE visualization of the *de novo* matrix, including 8,000,668 iSNVs, labelled with WHO lineage annotations. **F**: *de novo* iSNV matrix, as in **E**, but with *PNN_W HO_* labelling and the *PNN_W HO_* value at the top left. Where a lineage lacks a WHO designation, it is labelled as “Other,” and unassigned lineages are labelled as “Unassigned.”

This result is likely driven by lineage-specific mutations, herein referred to as consensus iSNVs, which are identified as having an Alternative Allele Frequency (*AAF*) over 75% and are usually part of consensus sequences (Ferreira et al. 2021; Murall et al. 2021; Thielen et al. 2021). These consensus iSNVs account for 3,634,563 iSNVs, averaging 28 per library (Table S1), aligning with the SARS-CoV-2 mutation rate reported by NextStrain (Hadfield et al. 2018) for this time period highlighting the reliability of these consensus iSNVs. PHATE representation of consensus iSNVs only again shows strong alignment with WHO lineages (Figure 3C), which is reflected in the high *PNN_W HO_* values in PHATE of 99.05% (Figure 3D), confirming these iSNVs largely drive the lineage-specific clustering observed in our raw iSNV dataset.

In contrast, we define putative *de novo* iSNVs (*AAF <* 0.75), representing emerging viral mutations within the host, totalling 8,000,668 in the raw dataset, averaging 62 per library (Table S1). The *de novo* iSNVs exhibit more heterogeneous clustering patterns (Fig. 3E) with a lower *PNN_W HO_* value of 54.83% (Figure 3F), suggesting a less pronounced lineage-based structure. However, the clustering patterns of the *de novo* iSNVs show a stronger alignment with lineage structure than expected by chance, with the baseline *PNN_W HO_* from random resampling at 32.82% for PHATE representation. This highlights the significance of the observed *PNN_W HO_* compared to the baseline value, suggesting a lineage-specific biological relevance in the emerging mutations. The controlled sub-sampling experiments (Figure S1, detailed in Methods section 4.5 and in supplementary information section 10.4) further support these observations, underscoring the distinct clustering behaviours of consensus and *de novo* iSNVs.

### 2.3 Resolving Artifacts in *de novo* iSNVs

Due to the geographic distribution of lineages, sequencing centers often sequence certain lineages more frequently than others, potentially leading to technical artifacts that affect lineage clustering in the *de novo* iSNV subset. This is confirmed by the clustering analysis of the 8,000,668 *de novo* iSNVs (Figure 4), where the PHATE representation showed significant sequencing center batch effects (Figure 4A), with a mean percentage of nearest neighbours from the same sequencing center (*PNN_SC_*) value of 62.31% (Figure 4B), greatly exceeding the baseline value of 27.53%, expected by chance. This result indicates that our set of *de novo* iSNVs likely contains sequencing artifacts. To filter out sequencing artifacts from the set of *de novo* iSNVs and refine the dataset, we used the strand bias metric *S* (see Methods, equation 1) and the Alternative Allele Frequency (*AAF*).

**Figure 4:**
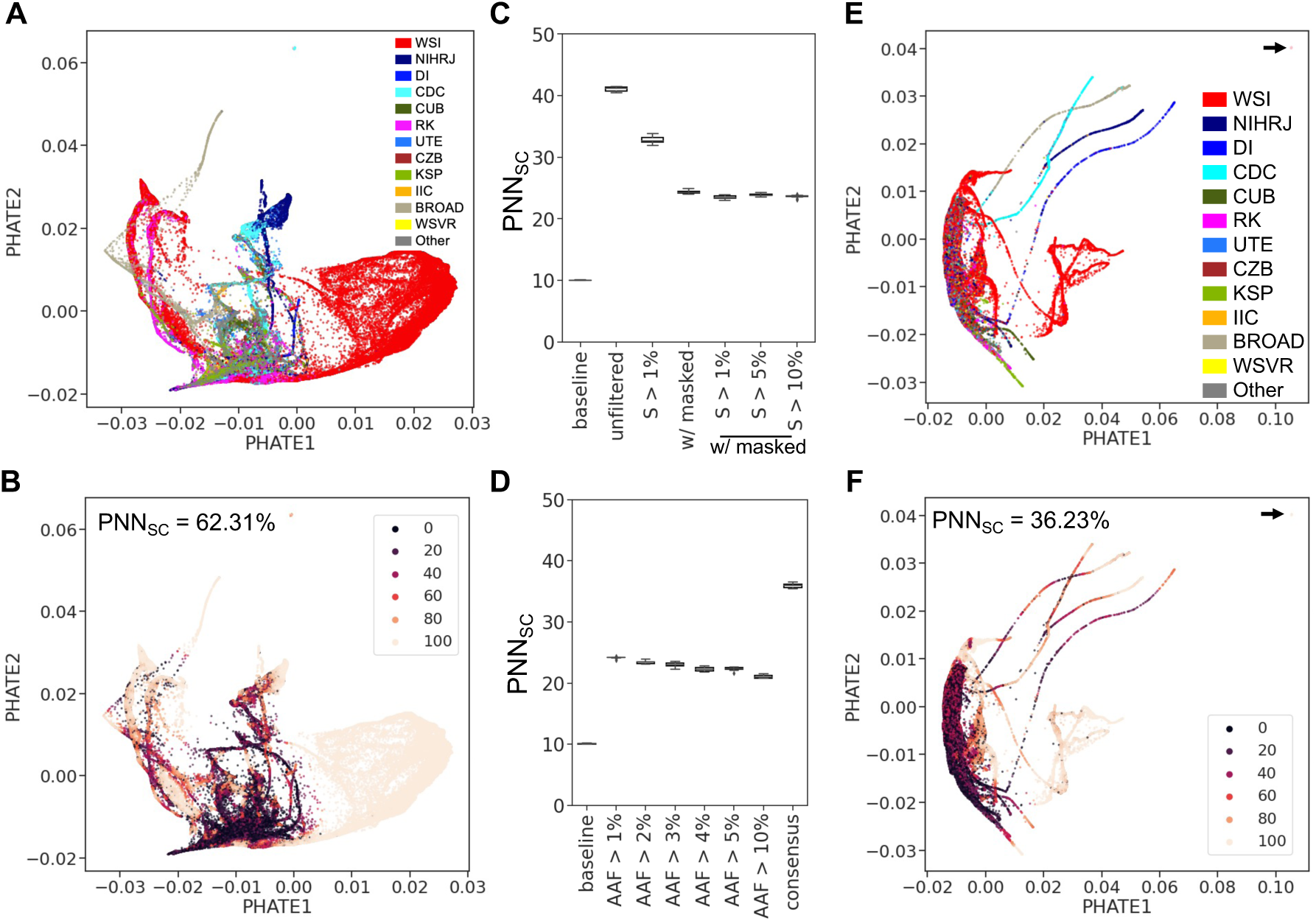
The Use of *S* and *AAF* Metrics Improves SARS-CoV-2 *de novo* iSNVs’ PHATE Structure by Mitigating Sequencing Center Batch Effects and Artifacts. **A**: PHATE visualization of the unfiltered *de novo* matrix containing 8,000,668 iSNVs, labelled by the libraries’ sequencing centers. **B**: PHATE visualization as in **A**, but with labels showing the percentage of *k*=100 nearest neighbours that share the same Sequencing Center (SC) annotation as the library itself and the total *PNN_SC_*value is displayed at the top left. **C** and **D**: Boxplots displaying *PNN_SC_* values for each PHATE visualization, derived from a sub-sampling controlled experiment across ten replicates (see method section 4.5). **C** shows *PNN_SC_*values across various *S* metric thresholds, and **D** presents *PNN_SC_* values across different *AAF* metric thresholds. **E**: PHATE visualization of the *de novo* matrix, filtered based on *S* and *AAF* thresholds, labelled by sequencing centers. **F**: PHATE visualization as in **E**, but with labels showing *PNN_SC_* values, and the total *PNN_SC_* value is displayed at the top left. In this representation sequencing centers with at least 1,000 libraries in our dataset are explicitly labelled with its sequencing center as follows: Welcome Sanger Institute (WSI), National Institute of Health DR. Ricardo Jorge (NIHRJ), Doherty Institute (DI), CDC-OAMD (CDC), Comenius University in Bratislava (CUB), Ravi Kant (RK), University of Tartu in Estonia (UTE), Chan Zuckerberg Biohub (CZB), Kwazulu-Natal Sequencing Platform (KSP), INAB Insitute in Certh (IIC), BROAD GCID (BROAD), Wales Specialist Virology Center (WSVR). While the remaining libraries were grouped under the “Other” label.

The PHATE visualization of the unfiltered *de novo* iSNVs prominently identifies the Wellcome Sanger Institute as a major cluster (Figure 4A) due to its significant representation of 75% in our library set. This underscores the potential impact of unbalanced sampling on cluster formation and potentially *PNN_SC_* values. To neutralize this imbalance, we designed a controlled sub-sampling experiment, evenly selecting 10,000 libraries from each of the top 10 sequencing centers based on library counts (see Method section 4.5), aiming to reduce the impact of sampling bias on the *PNN_SC_* values. We thus assessed the impact of filtering based on these two metrics, *S* and *AAF*, on the PHATE clustering structure measured with *PNN_SC_* using the controlled sub-sampling experiment to mitigate bias from uneven sampling across sequencing centers (Figure 4C, D).

To address the observed strand coverage unbalanced in our dataset (Figure 2C), we used the strand bias metric *S*, which assesses the likelihood of strand bias artifacts using the alternative allele’s strand coverage. Initially, filtering out iSNVs with *S <* 1% and 486 genomic positions showing recurrent strand bias across libraries (see supplementary information section 10.3) significantly lowers sequencing center-specific artifacts. This was reflected in the reduced *PNN_SC_* values (Figure 4C) in the controlled sub-sampling experiments. However, *PNN_SC_*values remained stable when the *S* threshold was increased beyond 1%, suggesting no further improvement based on this metric (Figure 4C).

Filtering based on allele frequency is a key metric in genomic studies. Some studies use a low threshold, which may result in the inclusion of erroneous intra-host mutations (Y. Wang et al. 2021; Armero et al. 2021; Popa et al. 2020; Tonkin-Hill et al. 2021; Lythgoe et al. 2021). In contrast, more stringent criteria could overlook the analysis of low-frequency, *de novo* intra-host mutations. Further refinement of our iSNV set based on the *AAF* metric led to an additional decrease in *PNN_SC_* values (Figure 4D), particularly when increasing the *AAF* threshold to 5%. Despite testing additional combinations of thresholds, the final *PNN_SC_* metric did not reach the baseline value of 10%, suggesting that the optimal threshold on the *AAF* metric is 5%.

Applying these optimal thresholds of 1% for *S* and 5% for *AAF* to filter out iSNVs, the *de novo* iSNV count dropped from 8,000,668 to 468,651, averaging six iSNVs per library (Table S1). This process notably decreased sequencing center batch effects in the PHATE representation (Figure 4E-F), resulting in a *PNN_SC_* value of 36.23%. While this value still exceeds the baseline of 27.69%, the reduction marks an improvement in minimizing batch effects. Additionally, the *PNN_SC_* value for lineage-defining consensus iSNVs also does not reach the baseline value (Figure 4D), implying that completely separating sequencing center influences from lineage-specific signatures might represent an intractable challenge.

### 2.4 Identifying Outliers and Center-Specific Patterns

In our analysis of the 468,651 filtered *de novo* iSNVs, there remain outlier clusters showing sequencing center homogeneity in the PHATE representation (Figure 4E-F). Notably, a small but distinct set of libraries forms an outlier cluster, markedly separated from other libraries in the PHATE representation (Figure 4E-F, indicated by an arrow). This observation suggests that specific libraries from the same sequencing centers potentially have an excess of shared iSNVs. We thus analyzed libraries’ intra-host mutational load, defined as the number of iSNVs in a library (see Method section 4.6). While most libraries in our dataset contain only one or two iSNVs (Figure S2), some exhibit a high intra-host mutational load, with tens of iSNVs per library.

To determine the optimal threshold for excess iSNVs in libraries, we computed the *PNN_SC_* value in PHATE representation after sequentially removing the top 1%, 5%, and 25% of the most mutated libraries (Figure S2A-B). Removing the top 1% of outliers impacted the *PPN_SC_* value the most, decreasing it by 2%. Additional exclusions, even down to only keeping libraries with one iSNV, did not further reduce the *PNN_SC_* value (Figure S2B, 50*^th^* percentile), underlining the impact of extreme outliers in the PHATE representation of the full dataset.

To ensure biologically relevant libraries are not excluded, we explored in-depth the patterns observed in the top 1% outlier libraries (1,270 outlier libraries) by computing the PHATE representation only on these libraries (Figure 5A, S3A). They strongly cluster by sequencing centers, indicating that their iSNVs are enriched for sequencing center-specific artifacts. In this PHATE representation of the outlier libraries, we note four main clusters (Figure S3). Cluster 1 is composed of 159 Doherty Institute libraries, corresponding to Australia’s first pandemic wave (March-August 2020) (Figure S3B). Cluster 2 comprises 104 libraries from Scilifelab Stockholm, collected at the end of the second pandemic wave. Cluster 3 includes 109 libraries from the Kwazulu-Natal Sequencing Platform, with collections from January to April 2021. Cluster 4 comprises 75 libraries from the Ravi Kant sequencing center, with a collection peak in May 2021.

**Figure 5:**
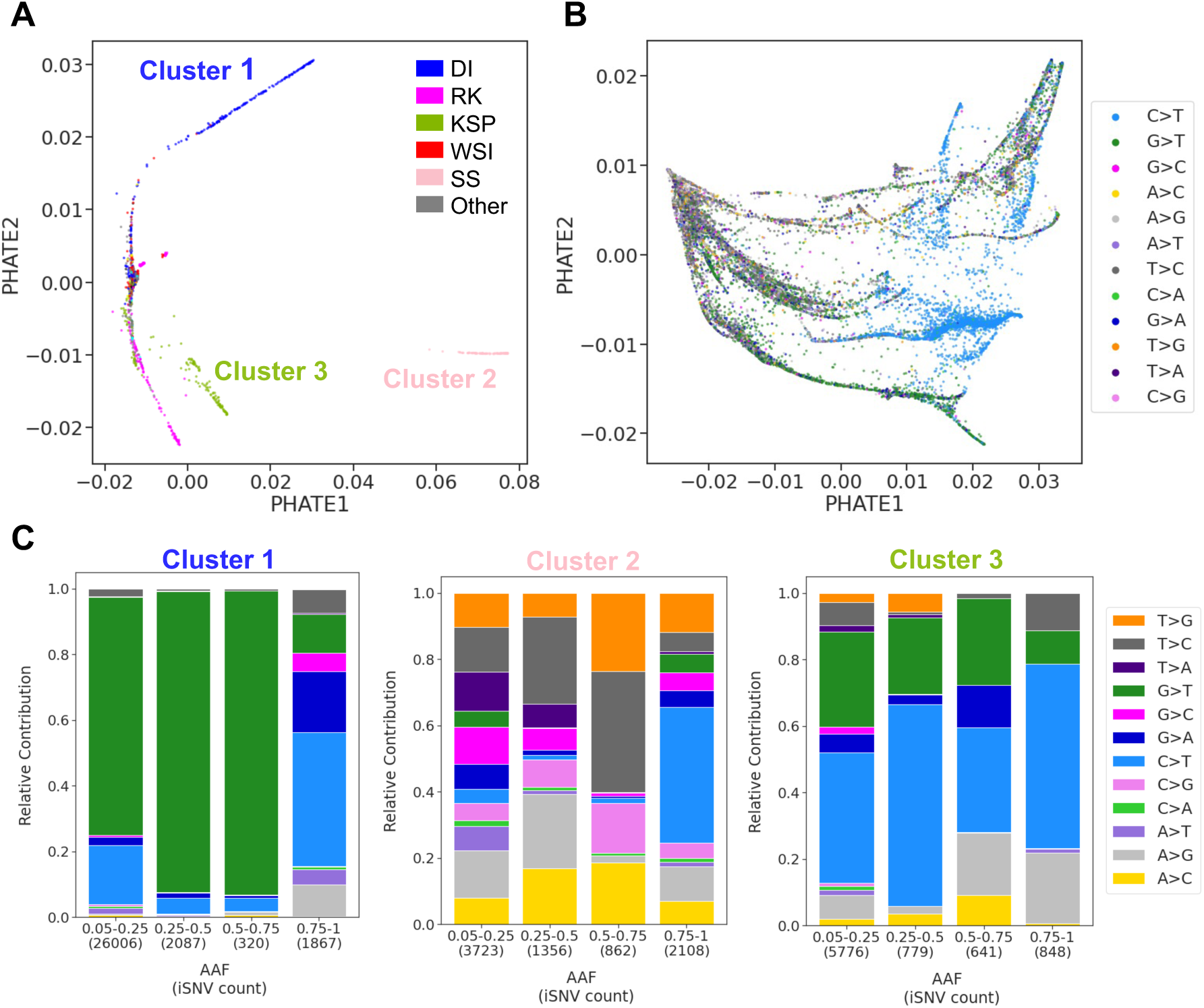
Unique Mutational Patterns in SARS-CoV-2 outlier libraries Tied to Sequencing Centers. **A**: PHATE visualization of outlier libraries, showcasing distinct clusters of SARS-CoV-2 libraries, each associated with specific sequencing centers. **B**: Displays the PHATE representation of genomic positions in outlier libraries, labelled with the most frequent substitution types observed across these libraries. **C**: Mutational patterns in iSNVs across the three distinct clusters in **A**, each associated with a specific sequencing center. **C** sequentially presents mutational patterns in iSNVs from Cluster 1 sequenced by the Doherty Institute, Cluster 2 associated with Scilifelab Stockholm, and Cluster 3 predominantly sequenced by the Kwazulu-Natal Sequencing Platform. The sequencing center’s labels are as follows: Welcome Sanger Institute (WSI), Doherty Institute (DI), Ravi Kant (RK), Kwazulu-Natal Sequencing Platform (KSP), and Scilifelab Stockholm (SS).

To detect the mutational patterns responsible for these effects, we computed PHATE representation by genomic position on these outliers libraries (Figure 5B), showing clustering of C*>*T and G*>*T mutations. This contrasts with non-outlier libraries, which do not show clear clustering based on substitution patterns (Figure 6A). Quantifying the proportion of iSNVs based on the substitution spectrum (see Methods section 4.7) revealed unique mutational signatures within each of the three main clusters with the most libraries (Figure 5C). Each cluster, associated with a specific sequencing center, exhibited mutational patterns distinct from those in non-outlier libraries (Figure 6A). Cluster 1 displays a prominent G*>*T pattern in *de novo* iSNVs, not seen in the consensus iSNVs from these same sequences. Interestingly, we identified 40 genomic positions with a *de novo* iSNV in at least 80% of the libraries in cluster 1. Cluster 2 libraries also displayed a unique mutational pattern in their iSNVs (Figure 5C, center), with T*>*G, T*>*C, A*>*G, and A*>*C as the predominant substitutions. These also diverged from their respective consensus iSNVs except for T*>*G. A notable 30 genomic positions have a *de novo* iSNV in at least 80% of the libraries in cluster 2. Lastly, cluster 3 libraries presented an excess of G*>*T and C*>*T that differed from their consensus iSNVs. In this cluster 3, 14 genomic positions have a *de novo* iSNV in at least 80% of the libraries.

**Figure 6:**
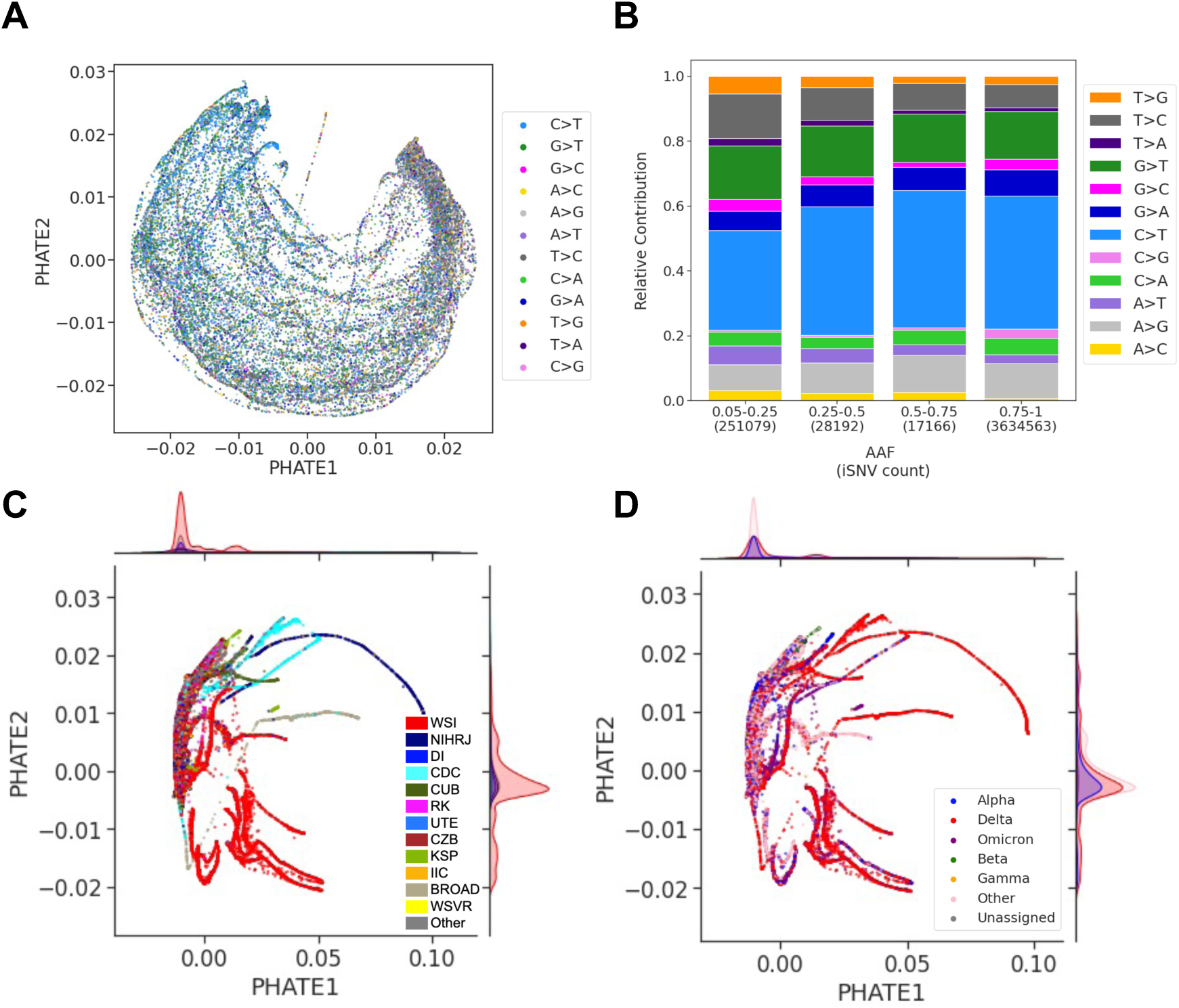
Attaining a Refined and Comprehensive Collection of SARS-CoV-2 Intra-host Sequencing Libraries and iSNVs via Meticulous Filtering. **A**: PHATE visualization of the refined library set, excluding outliers, with *de novo* iSNVs filtered based on *S* and *AAF* metrics. Each library is labelled by its sequencing center. **B**: Similar to **A**, but with labels showing the percentage of nearest neighbours (*PNN_SC_*) for each sequencing center and the total *PNN_SC_*value displayed at the top left. **C**: The total 296,437 *de novo* and consensus iSNVs, stratified by *AAF* and substitution types to reveal mutational biases. **D**: Presents a PHATE visualization of the transposed matrix for non-outlier libraries with filtered *de novo* iSNVs. Each point represents a genomic position of the SARS-CoV-2 genome, labelled by its most frequent substitution type across the libraries. Sequencing centers with at least 1,000 libraries are explicitly labelled with its sequencing center as follows: Welcome Sanger Institute (WSI), National Institute of Health DR. Ricardo Jorge (NIHRJ), Doherty Institute (DI), CDC-OAMD (CDC), Comenius University in Bratislava (CUB), Ravi Kant (RK), University of Tartu in Estonia (UTE), Chan Zuckerberg Biohub (CZB), Kwazulu-Natal Sequencing Platform (KSP), INAB Insitute in Certh (IIC), BROAD GCID (BROAD), Wales Specialist Virology Center (WSVR).

Overall, our outlier analysis revealed unique mutational patterns in *de novo* iSNVs across different sequencing centers associated with an excess of iSNVs, showing the influence of center-specific sequencing factors. These findings confirm the need to filter out the top 1% outlier libraries with a mutational load above 44 iSNVs in our library set. Our results also highlight the importance for sequencing centers to assess both the abundance of iSNVs and the presence of unique mutational patterns as key indicators for evaluating their sequencing processes.

### 2.5 Deriving a Final *de novo* iSNV Dataset

After our extensive curation, we kept 296,437 *de novo* iSNVs with *AAF >* 5% and *S >* 1%, from 72,470 non-outlier libraries with at least one iSNV, as our final curated dataset. The PHATE visualization of the 296,437 retained *de novo* iSNVs by genomic position display no clustering according to the mutational pattern, underscoring the optimal curation of the dataset (Figure 6A). Additionally, the substitution spectrum of this curated set shows a prevalence of C*>*T and G*>*T substitutions (Figure 6B), aligning with consensus iSNV patterns and known SARS-CoV-2 mutational trends (Moshiri et al. 2023; Fumagalli et al. 2023; Bloom et al. 2023; Saldivar-Espinoza et al. 2023).

The PHATE visualization by library (Figure 6C) shows greater sequencing center homogeneity compared to the initial representation of the raw iSNV data (Figure 4A). WHO lineage annotations of the same PHATE representation show similar lineage homogeneity (Figure 6D). Both sequencing center and WHO lineage annotations in the PHATE representation concentrate the majority of libraries into a single large cluster, as shown by the density plots. Despite the presence of sequencing center-specific clusters (Figure 6C), lineage-specific clustering is also noticeable (Figure 6D), suggesting that lineages from similar geographic regions may share iSNV generation processes, meriting further investigation. Nevertheless, the optimal refinement of the dataset is supported by a substantial decrease in the *PNN_SC_* value, from 62.31% to 33.26% (baseline value 26.29%).

## 3 Discussion

Emerging *de novo* mutations, or iSNVs, which occur during the intra-host phase of infection, are critical for understanding viral diversity and evolution. These mutations can be detected by analyzing sequencing libraries from infected hosts, although the sequencing process may introduce artifacts, resulting in false iSNV calls. To address this challenge, we present a comprehensive two-step workflow tailored for intra-host viral NGS analysis, specifically focusing on the SARS-CoV-2 RNA-seq libraries. It is specifically designed to robustly accommodate and correct for artifacts arising from the diverse sources present in our heterogeneous dataset, ensuring accurate detection of true iSNVs. First, we processed a large dataset of libraries with stringent whole genome quality control. Subsequently, we use these libraries for iSNV calling, employing specific quality metrics to differentiate putative iSNVs from artifacts. We also implemented dimensionality reduction techniques like PHATE and t-SNE to visualize and analyze library structures, enhancing our analysis with an explainability metric. Applying this workflow to a substantial SARS-CoV-2 dataset, we identified a set of emerging (*de novo*) iSNVs for studying intra-host viral evolution, differentiating them from consensus iSNVs using a 75% allele frequency threshold. This threshold is often used for its balance between detecting true positives and minimizing false positives, at the expense of intra-host diversity, by consensus callers (Ferreira et al. 2021; Murall et al. 2021; Thielen et al. 2021). Additionally, we tackled the challenge of distinguishing *de novo* iSNVs from similar-frequency artifacts using tailored quality metrics to establish appropriate thresholds for a given dataset, ensuring our process is rigorous and non-arbitrary.

Sequencing accuracy is influenced by multiple factors, including sample preparation, PCR amplification, and sequencing errors (Heguy et al. 2022; Zanini et al. 2017; McCrone et al. 2016; Grubaugh et al. 2019). This is especially the case when accurately detecting viral intrahost diversity (McCrone et al. 2016; Illingworth et al. 2017; Zanini et al. 2017). Mutations appearing on only one strand are likely due to amplification errors, as putative mutations would be present on both strands. Known as strand bias artifacts, they have been overlooked in the literature (Dinis et al. 2016; Illingworth et al. 2017; Zanini et al. 2017), but when addressed in recent studies, it is typically through applying a stringent filter that counts the appearances of an alternative allele on each strand (Sun et al. 2023; Xi et al. 2023; N’Guessan et al. 2023). However, this common filtering approach fails to account for the inherent imbalances in strand coverage frequently observed in targeted sequencing of SARS-CoV-2. This oversight can significantly increase the risk of false negatives, with the rate of missed variants varying unevenly across the genome. In response, our strand bias metric takes a different approach by assessing the distribution of each iSNV’s alternative allele across both strands, explicitly accounting for the imbalance in strand coverage observed in our SARS-CoV-2 NGS libraries. This approach avoids the bias of traditional methods that only retain genomic positions covered by both strands, a restriction that could impact about two-thirds of the genome (Figure 2C). Additionally, our strand bias metric, while similar to a published formula (McElroy et al. 2013), is tailored to a large viral NGS dataset. Interestingly, we highlight a set of genomic positions frequently identified as strand bias artifacts supported by our large and comprehensive dataset and see supplementary information section 10.3). By masking these positions, we noted a significant reduction in sequencing center batch effects, indicating that these positions may be specific to sequencing centers. Therefore, we highly recommend masking these positions to mitigate sequencing errors and erroneous data analysis and provide an efficient way to do so (Mostefai et al. 2024).

As intra-host viral genomic data grows in size and complexity (Chen et al. 2022; Smith et al. 2023), the challenge of managing these datasets increases. Dimensionality reduction methods are valuable for distilling this data into a more manageable form (Tapinos et al. 2019; Paradis 2022). However, interpreting these methods’ two-dimensional representations can be challenging due to unclear biological significance (Karim et al. 2022). In our workflow, we have incorporated PHATE and t-SNE alongside a metric that computes the percentage of nearest neighbours sharing the same annotation (e.g. sequencing center, WHO variant). This approach enhances the explainability of these techniques by highlighting relationships within specific groups of libraries in the representation, establishing a novel approach to analyzing high-dimensional viral sequence data. This methodology also facilitates the identification of optimal iSNV filtering thresholds, a critical aspect of sequencing data quality control. Implementing this approach allowed us to refine our quality metrics, resolve sequencing center batch effects, and improve the reliability of our iSNV dataset. Moreover, we have pioneered the use of PHATE in viral sequencing data analysis. We show that PHATE is especially effective at handling libraries with varying iSNV counts, unlike t-SNE, which is impacted by such libraries (see Figure S2 and supplementary information section 10.5). PHATE’s ability to accurately represent *de novo* mutations also demonstrates its potential for broader applications in areas requiring *de novo* mutation analysis, including the study of cancer clonal mutations (Muyas et al. 2023), evolutionary developmental biology (Short et al. 2018), and metagenomics (Keegan et al. 2016).

Despite PHATE’s ability to handle libraries with varying iSNV counts, outlier libraries containing a large number of iSNVs significantly skewed the PHATE clustering structure, highlighting a problematic aspect where a small subset disproportionately impacts the overall analysis. The significant influence of these outlier libraries was apparent in the unique C*>*T and G*>*T mutational patterns observed in PHATE’s genomic position representation within the outlier only libraries (Figure 5B), supporting the need to treat these libraries separately. Additionally, the strong clustering by sequencing center of the top 1% outlier libraries suggests that iSNVs within these libraries include sequencing artifacts specific to each center. This was confirmed by the distinct mutational patterns and the recurrence of genomic positions enriched for iSNVs within each outlier cluster, which, in turn, are associated with different sequencing centers. The unique mutational signatures identified within the outlier clusters also provide insight into the potential mechanisms of error introduction or bias in sequencing workflows. For instance, the G T substitution pattern seen in the Peter Doherty Institute libraries at the beginning of the pandemic (March to August 2020) may signal RNA degradation. Following the adoption of improved sample storage protocols, this Institute noted a reduction in the count of observed mutations (Peter Doherty Institute platform members, personal communication). This change in sequencing libraries’ quality emphasizes the necessity of ongoing collaboration between sequencing centers and data analysts to adapt practices and enhance sequencing data accuracy and reliability in real time.

Our approach, comprehensive as it is, faces some limitations. First, despite documented instances of mixed infections (Vatteroni et al. 2022; Rockett et al. 2022) where lineage-defining mutations appear at low frequencies, our current workflow is not designed to effectively capture these variations. In cases of mixed infections, iSNVs characterized as *de novo* under our definition may actually stem from the co-presence of two (or more) different strains within a host, as they would fall below our 75% threshold for emerging mutations. Therefore, particular care should be taken when analyzing datasets where a substantial proportion of samples could be mixed infections. In our analysis, we found little evidence of mixed infections, but it remains a possibility that some libraries—and consequently, iSNVs—could originate from such infections, though they would likely have been excluded during our outlier analysis. Furthermore, our workflow is currently not specifically designed to address the complexity associated with calling insertions and deletions (indels), which is an area for future development. In particular, a benchmark of indel detection tools for intra-host data should be conducted to enhance this aspect of viral genomic analysis. Bridging this gap poses a notable challenge and offers a valuable opportunity for methodological innovation. This workflow is designed specifically for Illumina sequencing data, favoured for its lower error rate (Fox et al. 2014), and is less suitable for nanopore sequencing (Cook et al. 2024; Fournelle et al. 2024). The latter’s high error rate of about 10% complicates the detection of low-frequency emerging mutations. Our dataset, while diverse, primarily consists of libraries from European and North American sources, mirroring the availability of publicly accessible sequencing data (Chen et al. 2022). This situation underscores the need for improved sequence sharing and support for sequencing capabilities in underserved regions. Additionally, our reliance on publicly available single instances of sequencing libraries leverages accessible data but complicates the confirmation of variant calls due to the absence of multiple replicates, as done previously. We address this by setting a minimum allele frequency threshold of 5%, higher than the typical Illumina error rate of 1% (Fox et al. 2014), aligning with the literature that advocates stricter thresholds for variant identification in the absence of replicates (Roder et al. 2023; Grubaugh et al. 2019).

Nonetheless, our workflow and dataset of high-quality intra-host iSNVs have proven instrumental in testing biological hypotheses and drawing conclusions on diverse areas of study. Published applications include uncovering immune evasion mechanisms in SARS-CoV-2 through sequence analysis and epitope mapping (N’Guessan et al. 2023), comparing intra-host viral evolution between immunosuppressed patients and the general population (Fournelle et al. 2024), and investigating intra-host mutations that influence epitope binding predictions (Caron et al. 2024). Additionally, this workflow and the identified set of *de novo* mutations open up new avenues for exploring hypotheses concerning viral intra-host diversity and evolution, providing a foundation for broader research initiatives in this field. For example, we observed that intra-host library clustering based on WHO variants persisted above baseline levels even after removing lineage-defining mutations. This leads us to hypothesize that lineage-defining genetic factors may contribute to the intra-host mutational patterns, suggesting a complex underlying mechanism of viral evolution within hosts. Our methodology has proven robust in detecting these subtle lineage characteristics despite variations in sample distribution, reinforcing the possibility of variant-specific effects on mutational events, a finding supported by a recent study (Bradley et al. 2024). This intriguing result warrants further investigation that could lead to the discovery of lineage dynamics and mutation impacts.

In conclusion, our robust viral intra-host processing and analysis workflow enhances the use of existing cross-sectional sequencing libraries and improves the accuracy and depth of viral genomic analyses. This advanced bioinformatics methodology is crucial for deepening our understanding of intra-host diversity and strengthening preparedness strategies for future pandemics, proving essential for responding effectively to other viruses in forthcoming outbreaks.

## 4 Methods

### 4.1 Data Selection and Library Pre-processing

We downloaded a set of SARS-CoV-2 Illumina amplicon paired-end sequencing libraries dataset from the first two years of the COVID-19 pandemic, ensuring a representative sampling across time and geographic locations. For each month from January 2020 to December 2021, sequencing libraries were randomly chosen based on availability in the National Center for Biotechnology Information (NCBI): up to 5,000 from the UK, up to 1,000 from the USA, and up to 2,000 from other global regions, totalling a potential 8,000 libraries monthly (Figure 2A). This yielded a total of 147,537 downloaded libraries (supplementary information section 10.1).

For each library, Illumina sequencing adapters and bad-quality reads (Phred score *<* 20) were trimmed from the sequencing reads using TrimGalore V.0.6.0 (https://github. com/FelixKrueger/TrimGalore). The trimmed libraries were mapped to the SARS-CoV-2 reference genome (NC045512.2) using BWA mem v.0.7.17-r1188 (Li et al. 2009), generating BAM files. Next, we used the iVar pipeline for primer trimming (Grubaugh et al. 2019), using the ARTIC Network V3, V4, and V4.1 amplicon designs, as the sequencing centers in our dataset predominantly use these three kits during the sampling period (https://github.com/artic-network/primer-schemes). We used the samtools mpileup (with specific parameters –Q 20 –q 0 –B –A –d 600000) (Danecek et al. 2021) to generate pileup files containing read information for each BAM file. To parse the pileup files and extract relevant data, we employed the publicly available script pileup2base (https://github.com/riverlee/pileup2base). We calculated the depth of coverage for each genomic position, which is the number of reads aligning to the position. The mean coverage across all libraries is 10446X, so we labelled any position with depth below 100X (1% of the mean) as low-quality. We calculated two metrics to evaluate each library’s quality (Figure 2B): (1) *C*, the mean coverage (the mean number of reads per position) and (2) *B*, the breadth of coverage (the number of genomic positions with a depth above 100X). We kept the libraries with *C >* 100X and *B >* 20,000 positions (representing two-thirds of the SARS-CoV-2 genome), yielding a total of 128,423 high-quality libraries. (see supplementary information section 10.1 for more details)

### 4.2 Consensus Sequences and Lineage Annotations

We obtained a consensus sequence for each of the 128,423 high-quality libraries using the iVar pipeline consensus calling tool (-q 20 –t 0.75 –m 20) (Grubaugh et al. 2019). These consensus sequences were annotated with Pango lineages using Pangolin 4.3 (Rambaut et al. 2020), which were next used to annotate with World Health Organization (WHO) lineages (Alpha, Delta, Omicron, Delta, Gamma, and Others) using a custom script. Sequences with no Pango lineage were annotated as ‘Unassigned’.

### 4.3 iSNV Calling and Encoding

We called iSNVs present in the 128,423 high-quality sequencing libraries (Figure 1B) after extracting genomic positions between positions 101 and 29,778, to exclude positions located at both ends of the genome that are generally of lower quality. For each library, we used pileup2base (Danecek et al. 2021) to obtain a base file, which contains, for the 29,678 positions, the counts for each nucleotide (A, T, C, G) separated according to amplicon direction (forward or reverse strand). Because we are focusing our analyses on single nucleotide substitutions, we ignored the last two columns of the base file that report the number of reads with indels. During this step, we kept only positions with a minimum coverage read depth of 100X.

We computed different iSNV metrics at the position level for each library using custom scripts. We define the alternative allele (*AA*) as the most frequent allele at a given position other than the reference allele. For each position and each library, we computed the Alternative Allele Frequency (*AAF*) as *AAF* = (*D_AA_*)*/D*, where *D* is the depth at the position studied and *D_AA_* is the depth for the alternative allele.

Due to the nature of targeted sequencing with amplicon design (Guo et al. 2012), it is possible that a single position in the genome may not be sequenced in a balanced manner between the forward and reverse directions. We thus compute the forward strand ratio as *FSR* = *D^f^ /D*, with *D^f^* the forward strand depth and *D* the total depth.

To evaluate if an alternative allele exhibits unbiased sampling across both strand directions, we used a binomial test. This test determines the probability of observing an allele predominantly on one strand, indicating a higher artifact likelihood. For the forward strand, let *Y* represent the expected count of reads bearing the alternative allele within the total forward strand reads, *D^f^*. Assuming *Y* follows a binomial distribution with a probability of success given by the *AAF*, the probability of observing at least 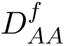 forward strand reads with the alternative allele (AA) is calculated as:

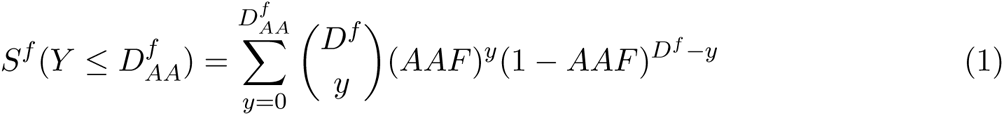

This same approach is applied to the reverse strand reads, *D^r^*, to calculate the likelihood of observing at least 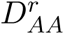 reverse strand reads with the alternative allele. Finally, to ensure a stringent evaluation, we take the minimum value of these calculated probabilities for both the forward and reverse strands. This minimum value serves as the Strand Bias Likelihood (*S*) metric for each iSNV, effectively quantifying the likelihood of no strand bias, and thus, a low value reflects the potential for the presence of an artifact.

The resulting iSNVs for each sequencing library are represented by their *AAF* given position *p* in a library *l* (*x_p,l_*) (Figure1B). This forms a matrix *X*, where the rows are our 128,423 high-quality sequencing libraries, and the columns are the genomic positions between 101 and 27778. We encode the initial unfiltered matrix *X* with *x_p,l_* = *AAF_p,l_* for all iSNVs from a given library *l*, and when an iSNV is filtered out based on thresholds for *AAF* and *S*, *x_p,l_* is set to 0.

### 4.4 Dimensionality Reduction and Clustering Evaluation

Given the high dimensionality of matrix *X*, we used dimensionality reduction methods to explore the underlying structure within the high-quality libraries in two dimensions (2D). We used incremental principal component analyses (PCA) for initialization and then obtained 2D representations of the PCA-transformed data using two different approaches: the widely-used t-distributed Stochastic Neighbor Embedding (t-SNE) (Tamazian et al. 2022; Maaten et al. 2008; Pedregosa et al. 2011) and the more recent heat diffusion for affinity-based transition embedding (PHATE) (Moon et al. 2019). We applied t-SNE with the Python library sklearn.manifold.T-SNE and PHATE with the PHATE Python library (Moon et al. 2019). The 2D embedding outputs from PHATE and t-SNE are visualized in scatter plots where each library is coloured either by sequencing center or WHO lineage annotation.

To measure the impact of specific subgroups of iSNVs on clustering structures based on either sequencing center (SC) or WHO lineage labels, we used a k-nearest neighbour (kNN) approach, using *k*=100. This value of *k* is selected to simplify interpretation as a percentage during neighbour selection and reflects the large number of libraries in our dataset. For each library *l* within a representation, we identify the 100 nearest libraries *NN* (*l*) using the sklearn.neighbors Python package (Pedregosa et al. 2011). We then calculate *NN_z_*(*l*), the count of nearest neighbours sharing the same *z* label as library *l* (where *z* is either *WHO* for lineage or *SC* for sequencing center), and compute the percentage of nearest neighbours with matching labels. For each representation, we derive a final *PNN_z_* as the mean percentage of nearest neighbours with matching labels across all libraries. A higher *PNN_z_* indicates that label *z* describes the data’s clustering structure. We also generate a baseline *PNN_z_*, representing expected chance levels by randomly shuffling labels *z* before calculation. This baseline acts as a standard for assessing the significance of observed patterns, emphasizing the delta between observed and baseline *PNN_z_* over the choice of *k* value.

### 4.5 Experimental Design to Mitigate Sampling Biases

We designed a controlled sub-sampling experiment by randomly sub-sampling libraries for each WHO or sequencing center annotation to address the impact of biases stemming from unbalanced sampling. To evaluate the influence of iSNVs on WHO patterns, we iteratively sampled 1000 library rows for each of the Alpha, Beta, Delta, and Omicron variants from the data matrix *X* ten times, generating replicates. This process resulted in ten matrices, each comprising 4,000 rows. To investigate the effect of iSNVs on sequencing center patterns, we used a similar approach, randomly selecting 1000 library rows. However, in this case, we randomly sampled 1000 library rows from our dataset’s top 10 most frequent sequencing centers (Table S3). This process resulted in ten matrices as replicates, each comprising 10,000 rows. Within each matrix, *x_p,l_*values were set to 0 based on various *AAF* and *S* thresholds cutoffs. After these two steps, we applied the same method as in Data Visualization to generate PHATE and t-SNE visualization of the matrices. Subsequently, we quantified the clustering structure of t-SNE and PHATE to derive a *PNN_z_* value for each visualization. Specifically, a high value of *PNN_S_C*, indicating clustering primarily by sequencing center, would suggest a dataset enriched for artifacts. Conversely, a high value of *PNN_W_ HO*, signifying clustering primarily by WHO lineage annotations, would suggest a more biologically relevant dataset.

### 4.6 Mutational Load

The mutational load for each library was calculated as the total count of distinct iSNVs identified regardless of allele frequencies. Per library, mutational loads were visualized using histograms to illustrate the distribution of mutational loads across the dataset. We categorized the libraries into different percentiles based on their mutational load, identifying those with higher or lower numbers of iSNVs.

### 4.7 Substitution Spectrum Analysis

To assess the mutational landscape and identify specific patterns that may indicate underlying mutational mechanisms or biases in the dataset, we looked at the substitution patterns within iSNVs’ different *AAF* frequencies. First, we categorized intra-host iSNVs into four *AAF* bins, as follows: 5% to 25%, 25% to 50%, 50% to 75%, and 75% to 100%. This categorization was based on the evidence for an alternative allele present in the iSNVs. Next, within each *AAF* bin, we classified each iSNV in terms of its ancestral allele and alternative allele to obtain 12 categories of substitution types. These are A*>*G, A*>*C, A*>*T, C*>*A, C*>*G, C*>*T, G*>*A, G*>*C, G*>*T, T*>*A, T*>*C, and T*>*G. This allowed us to analyze the relative contribution of each substitution type within each *AAF* range.

## 5 Data and Code Access

The processing workflow’s code can be found here: https://github.com/HussinLab/IntraHost_ Covid_Pipeline.git. NCBI accession IDs utilized in this study and the high-quality iSNVs identified within each sequencing library are accessible through a Mendeley data repository (Mostefai et al. 2024, https://doi.org/10.17632/8nvgtrkzdm.1). We also provide the list of recommended 477 genomic positions to mask in the same data repository (see supplementary information section 10.3).

## 7 Acknowledgments

The study was conducted in accordance with the Declaration of Helsinki and approved by the Ethics Committee of the Montreal Heart Institute (protocol code 2021-2868 and date of approval 23 July 2021). We express our gratitude to the members of Julie Hussin’s group for their valuable discussions. The successful completion of this work was made possible by the computational resources provided by the Digital Research Alliance of Canada clusters Narval and Beluga. We also thank sequencing library submitters who made data available on NCBI, who are listed in the available iSNV table (Mostefai et al. 2024). We especially extend our appreciation to the Peter Doherty Institute for their insightful communication regarding the distinctive mutational patterns observed within our dataset.

## 8 Authors’ Contributions

FM: conception, data acquisition, data pre-processing, data analysis, figure development, and manuscript drafting. RP and JCG: data acquisition, data pre-processing, data analysis, and manuscript critical revision and editing. JGH: funding, conception, supervision, and co-drafting of the manuscript.

## 9 Supplemental Tables and Figures

**Table S1:** iSNV Count, Library Count, and Mutational Load.

|  | iSNV Count | Library Count | Mutational Load |
| --- | --- | --- | --- |
| Total iSNVs | 11,635,231 | 128,443 | 91 |
| Consensus iSNVs | 3,634,563 | 128,323 | 28 |
| *De novo* iSNVs | 8,000,668 | 128,352 | 62 |
| S > 1% filtered *de novo* iSNVs | 6,508,783 | 127,941 | 51 |
| Masked *de novo* iSNVs | 5,805,486 | 125,382 | 46 |
| AAF > 5% filtered *de novo* iSNVs | 468,651 | 73,729 | 6 |
| Non-Outlier libraries *de novo* iSNVs | 296,437 | 72,470 | 4 |

**Table S2:**
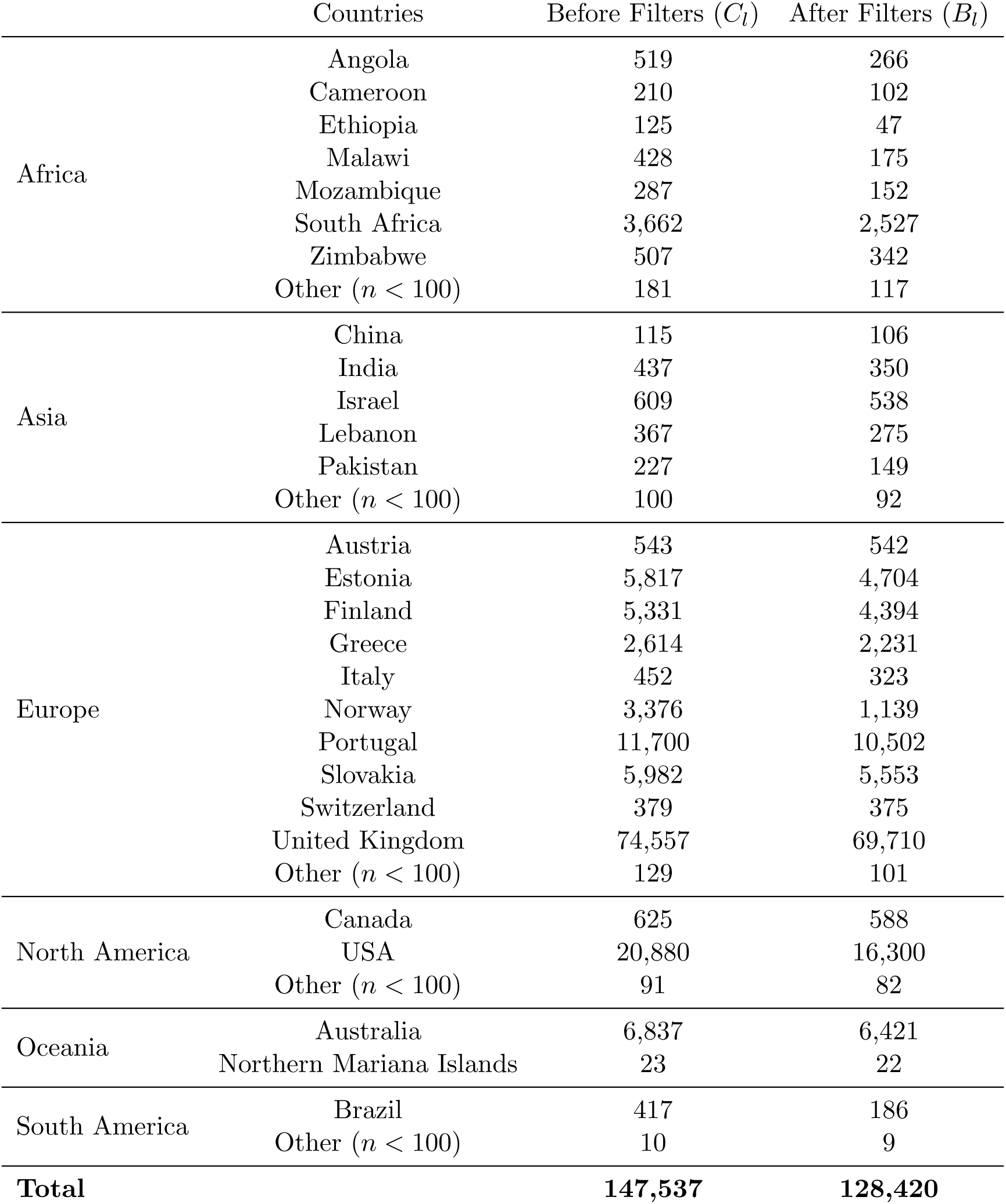
Per Country Sequencing Libraries’ Counts Before and After Coverage Filters.

| | Countries | Before Filters ($C_l$) | After Filters ($B_l$) |
| --- | --- | --- | --- |
| Africa | Angola | 519 | 266 |
|  | Cameroon | 210 | 102 |
|  | Ethiopia | 125 | 47 |
|  | Malawi | 428 | 175 |
|  | Mozambique | 287 | 152 |
|  | South Africa | 3,662 | 2,527 |
|  | Zimbabwe | 507 | 342 |
| | Other ($n < 100$) | 181 | 117 |
| Asia | China | 115 | 106 |
|  | India | 437 | 350 |
|  | Israel | 609 | 538 |
|  | Lebanon | 367 | 275 |
|  | Pakistan | 227 | 149 |
| | Other ($n < 100$) | 100 | 92 |
| Europe | Austria | 543 | 542 |
|  | Estonia | 5,817 | 4,704 |
|  | Finland | 5,331 | 4,394 |
|  | Greece | 2,614 | 2,231 |
|  | Italy | 452 | 323 |
|  | Norway | 3,376 | 1,139 |
|  | Portugal | 11,700 | 10,502 |
|  | Slovakia | 5,982 | 5,553 |
|  | Switzerland | 379 | 375 |
|  | United Kingdom | 74,557 | 69,710 |
| | Other ($n < 100$) | 129 | 101 |
| North America | Canada | 625 | 588 |
|  | USA | 20,880 | 16,300 |
| | Other ($n < 100$) | 91 | 82 |
| Oceania | Australia | 6,837 | 6,421 |
|  | Northern Mariana Islands | 23 | 22 |
| South America | Brazil | 417 | 186 |
| | Other ($n < 100$) | 10 | 9 |
| **Total** |  | **147,537** | **128,420** |

**Table S3:** Per Sequencing Center Librarie’s Counts Before and After Coverage Filters.

| Sequencing Center | Before Filters ($C_l$) | After Filters ($B_l$) |
| --- | --- | --- |
| Wellcome Sanger Institute | 69,676 | 65,316 |
| National Institute Of Health DR. Ricardo Jorge | 11,700 | 10,502 |
| Doherty Institute | 6,699 | 6,306 |
| CDC-OAMD | 6,431 | 5,040 |
| Ravi Kant | 5,331 | 4,394 |
| Kwazulu-Natal Sequencing Platform | 4,711 | 2,827 |
| Comenius University in Bratislava | 4,648 | 4,458 |
| University Of Tartu, Estonia | 4,104 | 3,351 |
| Norwegian Institute of Public Health (NIPH) | 3,376 | 1,139 |
| INAB Institute, Certh | 2,614 | 2,231 |
| BROAD, GCID | 2,334 | 1,948 |
| Chan Zuckerberg Biohub | 1,971 | 1,850 |
| Quadram Institute Bioscience | 1,859 | 1,257 |
| UPHL ID | 1,853 | 1,062 |
| Institute of Biomedicine and Translational Medicine | 1,713 | 1,353 |
| Wales Specialist Virology Centre | 1,507 | 1,485 |
| Chan Zuckerberg Biohub | 1,364 | 1,229 |
| Public Health Authority of the Slovak Republic | 1,349 | 1,106 |
| TX-SARS-COV-2 | 1,324 | 710 |
| Public Health England (Colindale) | 1,142 | 1,123 |
| DCLS-NGS | 1,062 | 492 |
| California Department of Public Health | 982 | 900 |
| Liverpool Clinical Laboratories | 825 | 741 |
| CanCOGeN CPHLN | 612 | 579 |
| Tel Aviv University | 609 | 538 |
| NYC SARS-COV-2 | 562 | 493 |
| CeMM | 543 | 542 |
| New Mexico Department of Health Scientific Laboratory | 539 | 503 |
| Colorado Department of Public Health and Environment | 509 | 463 |
| Gujarat Biotechnology Research Centre | 436 | 349 |
| West of Scotland Specialist Virology Centre, NHS GG | 362 | 362 |
| University Hospital of Basel | 336 | 336 |
| Delaware Public Health Lab | 327 | 299 |
| University of Kwazulu-Natal | 277 | 180 |
| Network for Genomic Surveillance in South Africa | 275 | 274 |
| CDC-PDD | 253 | 222 |
| Utah Public Health lab | 233 | 169 |
| Kwazulu-Natal Research and Sequencing Platform | 229 | 137 |
| UMIGS | 223 | 209 |
| University of Verona | 220 | 188 |
| SEARCH | 211 | 191 |
| Hospital Israelita Albert Einstein | 208 | 0 |
| SciLifeLab Stockholm | 164 | 133 |
| Institute of Clinical Pathology and Medical Research | 138 | 135 |
| LNCC | 119 | 110 |
| Ruijin Hospital, Shanghai Jiao Tong University of Medicine | 112 | 103 |
| Centers for Disease Control and Prevention | 100 | 95 |

**Figure S1:**
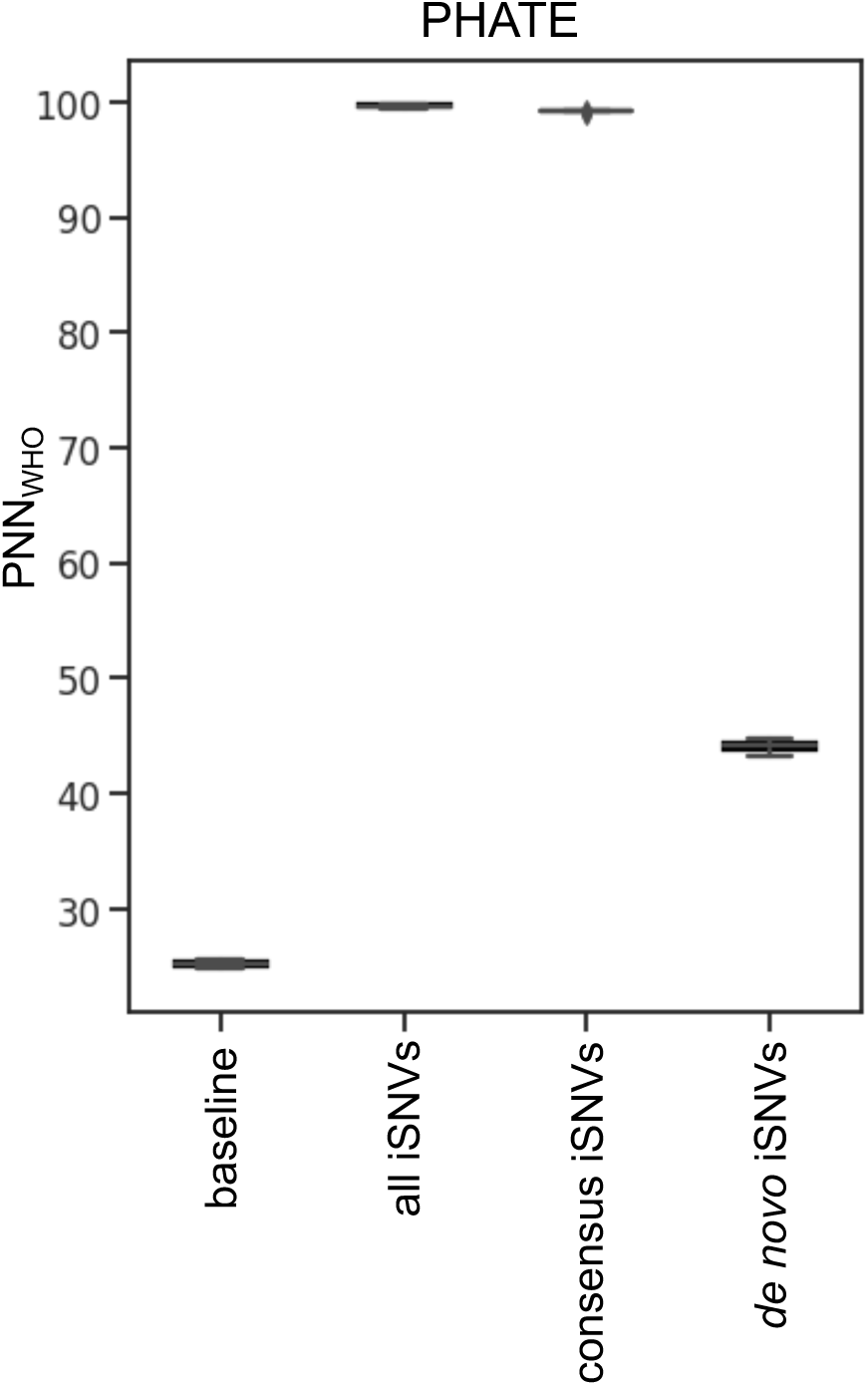
Unveiling WHO Lineage Patterns in SARS-CoV-2 iSNVs with PHATE Visualizations and *PNN_W HO_* Metric. Boxplots show the distribution of the mean percentage of nearest neighbours (*PNN_W HO_*) from the same WHO lineage annotation across libraries for each PHATE (**A**) visualization across the ten replicates from the sub-sampling controlled experiment (see method section 4.5). Before computing *PNN_W HO_*, PHATE visualizations were generated on matrices containing a consistent sampling of 4,000 libraries from each of Alpha, Delta, Omicron, and Beta WHO annotated lineages. For PHATE, the first boxplot represents the expected *PNN_W HO_* values by chance, followed by all iSNVs, consensus only iSNVs, and *de novo* only iSNVs. The number of nearest neighbours used in this experiment is *k*=40.

**Figure S2:**
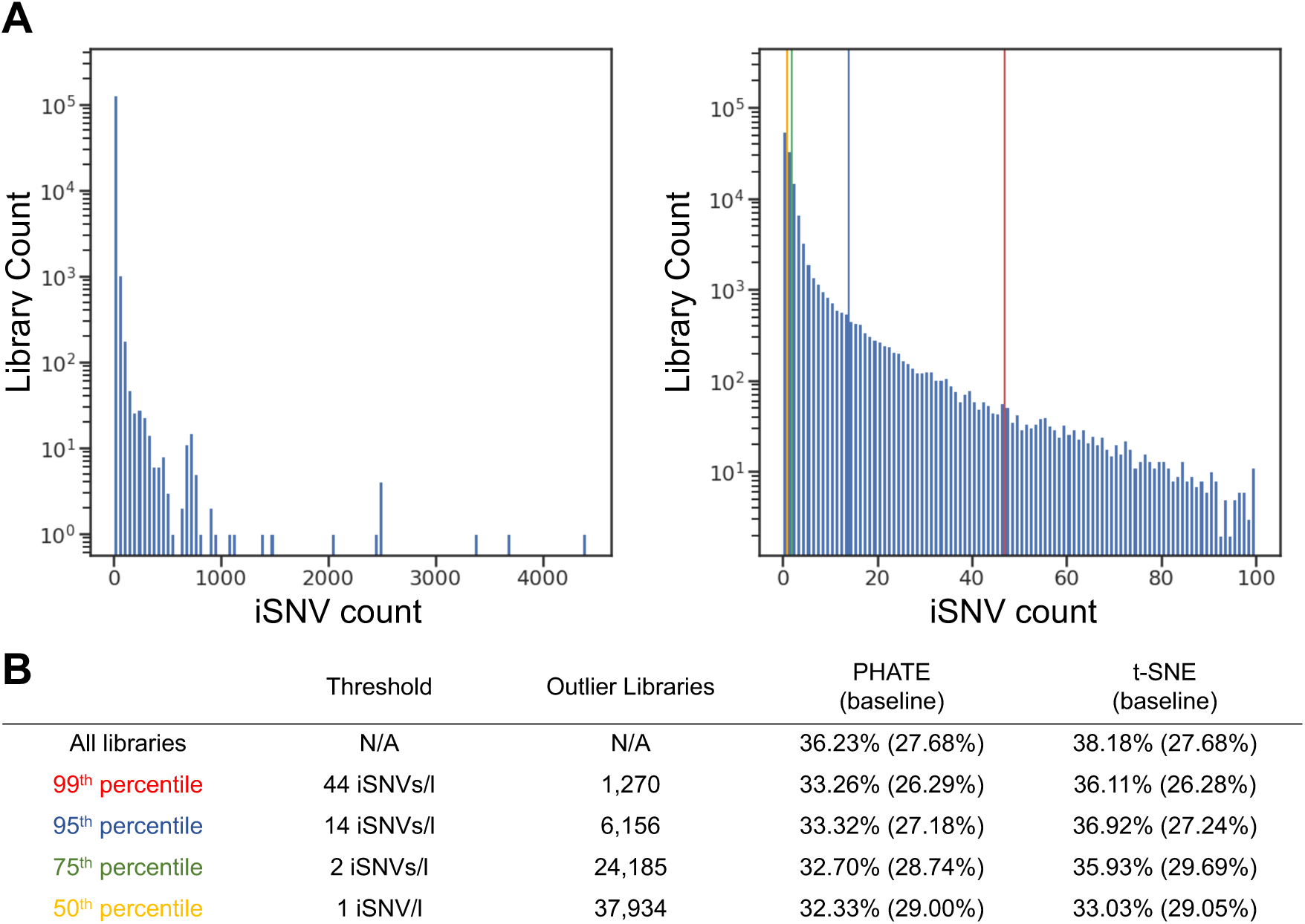
Analysis of Libraries’ Mutational Load, which is the Number of iSNVs per Library. **A** The mutational load distribution across all libraries shows the variability in the number of iSNVs per library. **B** zoomed-in view of this distribution, focusing on libraries with up to 100 iSNVs. This view includes vertical lines to delineate various distribution percentiles. **C** A table summarizing the relationship between different outlier detection thresholds and their impact on library clustering structure on the PHATE visualizations. The table shows thresholds defined by the number of iSNVs per library, ranging from the 99th percentile (44 iSNVs/library) to the 50th percentile (1 iSNV/library). For each threshold, the table indicates the number of libraries classified as outliers and the corresponding percentage of nearest neighbours from the same WHO lineage (*PNN_W HO_*), alongside the expected by chance *PNN_W HO_* value.

**Figure S3:**
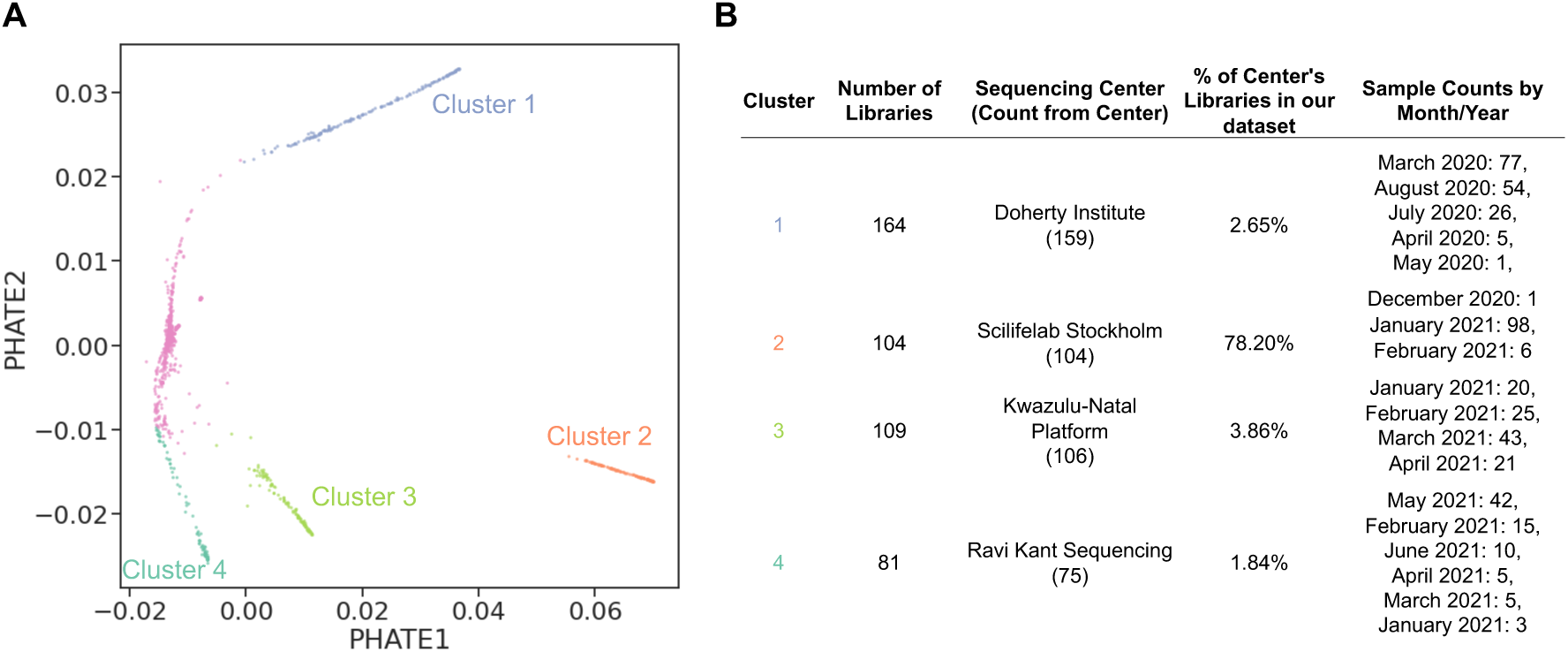
**A**: PHATE on the top 1% outlier libraries with the most iSNV count. The clusters on this PHATE representation were defined using K-means applied to the PHATE object. **B**: Table representing per cluster library information.

## 10 Supplemental Information

### 10.1 Details on Downloading SARS-CoV-2 Genomic Libraries from NCBI

A total of 147,537 SARS-CoV-2 Illumina amplicon paired-end sequencing reads were downloaded from NCBI, as follows: 51,837 Illumina SARS-CoV-2 sequencing libraries were downloaded from the NCBI database on June 4th, 2021, and another 95,700 Illumina sequencing libraries were downloaded on February 12th, 2022. January and February 2020 were severely underrepresented compared to the other months (Figure 2A). Most downloaded sequences originated from Europe, constituting 75% of the dataset. Among the European sequences, 63% were obtained from the Wellcome Sanger Institute sequencing center, UK (Table S3), indicating their significant contribution to the global sequencing efforts. Furthermore, a notable number of downloaded libraries came from The Peter Doherty sequencing center, Australia, between January and October 2020 (16% of the total libraries Table S3) as they led the sequencing effort during that time in that region. Additionally, the dataset was enriched with samples sequenced by North American sequencing centers, accounting for 15% of the downloaded sequences (Tables S2 and S3). The underrepresentation of samples from January and February 2020 reflects a limitation in the available data during the initial stages of the pandemic. However, despite the initial disparities in data collection, which reflect the current practical challenges faced by the scientific community (Chen et al. 2022), this dataset remains highly informative, successfully capturing the global diversity of SARS-CoV-2 throughout the later months of 2020 and extending into 2021.

Out of the total libraries downloaded, 134,879 had a mean coverage *C* above 100, and a total of 138,723 libraries had a breadth of coverage *B* above 10,000, meaning that at least 10,000 genomic positions were covered at a depth of 100X or higher (Figure 2). The intersection of both filters allowed us to keep 128,423 high-quality sequencing libraries for further analysis. The distributions of the breadth of coverage and mean depth show heterogeneity in the coverage of the downloaded sequencing libraries. We also note the grouping of some sequencing centers (e.g. Welcome Sanger Institute in red) and not others (e.g. the CDC’s Office of Advanced Molecular Detection – CDC-OAMD), displaying a heterogeneity across sequencing centers and within sequencing centers. Because we downloaded a representative sampling of the available data on the NCBI database, this coverage distribution likely represents the coverage heterogeneity of the available data on NCBI.

### 10.2 Strand Coverage Across the Genome

We evaluated the variation in strand coverage along the genome in our dataset using the Forward Strand Ratio (FSR), which revealed a highly unbalanced distribution across the virus sequence (Figure 2C). Only 31% of the viral genome in our dataset has a balanced coverage from the forward and reverse read strands. Specifically, 40% of the genome is covered by the plus strand, which is the number of genomic positions of the genome with an average forward strand ratio above 90%. In contrast, 29% of the genome is covered by the minus strand, with an average minus strand ratio above 90%. Thus, strand bias statistics in SARS-CoV-2 genomes need to consider strand coverage when evaluating if a *de novo* iSNVs is a stand bias artifact, which motivates the development of our strand bias likelihood metric *S*.

### 10.3 Recurrent Strand Bias Artifacts

To better characterize strand bias artifacts, we analyzed a total of 1,491,885 intra-host single nucleotide variants (iSNVs) identified as potential strand bias artifacts, with a likelihood of no strand bias below 1% (*S*¡0.01). We first examined their alternative allele frequency (*AAF*) distribution. The *AAF* distribution of these excluded iSNVs does not differ significantly from that of the other iSNVs, suggesting that strand bias artifacts can happen across a spectrum of intra-host frequencies. This confirms that filtering based solely on *AAF* is insufficient to eliminate strand bias artifacts.

Several genomic positions were found to be recurrent within these putative strand bias artifacts. We computed the expected number of libraries with strand bias artifacts at a given position, which has a mean of 4 and a 99th percentile of 68 libraries. We identified 486 genomic positions that have a strand bias artifact reported in more than 68 libraries, labelling them as recurrent strand bias artifacts, which we masked in our analyses across all libraries. To ensure the robustness of iSNV analyses and to prevent the inclusion of recurrent spurious iSNVs, we recommend evaluating and possibly masking these genomic positions in future SARS-CoV-2 intra-host studies.

### 10.4 Sub-sampling experiments to balance WHO variants

In our dataset, Alpha and Delta are overrepresented compared to other SARS-CoV-2 variants, which may cause biases in the analysis results since unbalanced sampling can influence cluster formation and *PNN_W HO_* values (see, for example, Figure 3A, which distinctly marks Alpha and Delta as dominant clusters). To address this, we conducted controlled sub-sampling experiments, selecting 1,000 libraries each from the Alpha, Beta, Delta, and Omicron variants (see Method section 4.5), aiming to mitigate variant sampling bias on *PNN_W HO_* values in the PHATE representation of iSNV subsets. We evaluated the clustering by WHO lineage across three iSNV sets: unfiltered raw iSNVs, consensus iSNVs, and *de novo* iSNVs (Figure S1). The raw and consensus iSNV datasets show high *PNN_W HO_* values, indicating a strong lineage-specific signature, primarily driven by frequent lineage-defining mutations, even when samples per WHO variant are balanced. Conversely, *de novo* iSNVs exhibit lower *PNN_W HO_* values, indicating a subtler lineage-based structure but still above baseline, underscoring the lineage-specific biological significance of emerging mutations. These controlled subsampling experiments thus replicate our main findings with the full dataset (Figure 4). Therefore, the lineage-specific signatures observed in our study are not a result of the uneven sampling of WHO variants.

### 10.5 t-SNE Results Are Comparable to PHATE

In this section, we present results from t-SNE (t-Distributed Stochastic Neighbor Embedding) analysis of SARS-CoV-2 genomic data, complementing the PHATE results found in the result section (2). The method t-SNE is a machine learning algorithm used for dimensionality reduction, offering an alternative approach to PHATE.

The t-SNE representation of the 128,423 high-quality sequencing libraries reveals distinct clusters by WHO lineage for both raw and consensus iSNV subsets, consistent with PHATE’s findings. For raw iSNVs, the Proportion of Nearest Neighbors (*PNN_W HO_*) for t-SNE is 99.43%, closely aligned with PHATE’s 98.39%. Similarly, for consensus iSNVs, tSNE’s *PNN_W HO_* of 99.05% parallels PHATE’s 99.37%, highlighting both methods’ consistent ability to identify lineage-specific mutations across the iSNV sets. Conversely, *de novo* iSNVs (representing emerging mutations within the host) show less pronounced lineage-specific than consensus iSNVs clustering in t-SNE representation, with a *PNN_W HO_* value of 59.37%. This suggests a deviation from the strong lineage alignment observed in raw and consensus iSNVs, indicating that while *de novo* iSNVs still correlate with lineage structure more than baseline, the association is less direct. The structure observed in *de novo* iSNVs through t-SNE complements PHATE’s analysis, demonstrating consistent underlying data patterns regardless of the representation method used.

Using the 8,000,668 unfiltered *de novo* iSNVs, both t-SNE and PHATE visualizations revealed significant sequencing center batch effects, with t-SNE showing slightly higher *PNN_SC_* values (66.50%) compared to PHATE (62.31%). This indicates that both dimensionality reduction techniques captured the influence of sequencing center-specific artifacts within the *de novo* iSNV dataset. Efforts to refine the dataset and mitigate these artifacts involved applying thresholds on the strand bias metric (*S*) and the Alternative Allele Frequency (*AAF*). These measures effectively reduced sequencing center-specific artifacts, as evidenced by decreased *PNN_SC_* values in both visualization methods after applying the filters, with the t-SNE value (38.18%) slightly higher than PHATE (36.23%). Applying the filters effectively reduced sequencing center-specific artifacts, as evidenced by decreased *PNN_SC_* values in both representation methods.

Similarly to PHATE, we also computed the *PNN_SC_* values in t-SNE representation after sequentially removing the top 1%, 5%, and 25% of the libraries with the most iSNV counts (Figure S2B). As opposed to PHATE, the *PNN_SC_* value of t-SNE did not drastically decrease after the removal of the top 1% of our outliers. However, the *PNN_SC_* values for both t-SNE and PHATE only met after the exclusion of more libraries down to only keeping libraries with one iSNV (Figure S2B, 50*^th^* percentile), underlining the stronger impact of outlier libraries on t-SNE compared to PHATE.

Similar to the approach used with PHATE, we calculated the *PNN_SC_*values for t-SNE after removing the top 1%, 5%, and 25% of libraries based on iSNV counts (Figure S2B). Unlike PHATE, the *PNN_SC_* for t-SNE did not significantly decrease with the removal of the top 1% of libraries. Both t-SNE and PHATE *PNN_SC_* values converged after removing more libraries, ultimately comparable for their *PNN_SC_* values only when retaining those with a single iSNV (Figure S2B, 50*^th^* percentile). This indicates that t-SNE is more susceptible to bias from outlier libraries compared to PHATE.

This overall consistency between dimensionality reduction methods serves as compelling evidence that the data’s underlying structure is method-independent, suggesting that both methods could be reliably applied to similar datasets to help inform future pre-processing strategies in viral genomics. This alignment helps validate our pre-processing strategies in viral genomics, demonstrating the robustness of our observations and the general applicability of these techniques to analyze viral genomic data.

